# RNA splicing is required for timely completion of abscission and is modulated by the abscission checkpoint

**DOI:** 10.1101/2025.11.09.687034

**Authors:** Genevieve C. Couldwell, Jessica N. Vincent-Mueller, Douglas R. Mackay, Christopher C. Jensen, Chris J. Stubben, Li Li, Aik Choon Tan, Wesley I. Sundquist, Katharine S. Ullman

## Abstract

The abscission checkpoint pauses the progression of cytokinetic abscission at the intercellular bridge when various checkpoint-sustaining errors are sensed. A unique cytoplasmic organelle, the abscission checkpoint body (ACB), forms when the abscission checkpoint is active. This organelle houses proteins involved in abscission and RNA splicing factors. Here, we investigate the role of RNA splicing factors in abscission checkpoint maintenance. A specific subset of RNA splicing factors localizes to ACBs, while others move back into the nucleus. The key RNA splicing factor SF3b1 is aberrantly localized in the cytoplasm and its catalytic activity limited when ACBs are present. Efficient abscission was found to rely on RNA splicing, and we have identified patterns in gene expression and RNA splicing that occur at this time in cell division. Finally, we found a unique RNA landscape associated with the formation of ACBs due to a specific set of changes in RNA expression and alternative splicing, revealing a previously unrecognized level of regulation imparted by abscission checkpoint activity.

## INTRODUCTION

Elucidating the steps in cell division is of critical importance to understanding developmental biology and tumorigenesis, and fundamental insights have been gained from the study of mitotic mechanisms and the transitions into and out of S phase [1–3]. The events of cytokinesis, though less thoroughly explored, are emerging as a critical regulatory nexus in development [4–12] and cancer biology ([13], reviewed in [14, 15]). The final cell cycle checkpoint occurs at cytokinetic abscission and is referred to as the abscission or NoCut checkpoint [16, 17]. This checkpoint prevents DNA damage and resolves mitotic errors before daughter cells separate. Abscission checkpoint activity can be sustained by four known stressors: DNA traversing the abscission site, heightened intercellular tension between the dividing daughter cells, replication stress earlier in the cell cycle, and depletion of specific nuclear pore proteins [18–21]. However, molecular mechanisms that link these physical inputs to NoCut signaling and abscission delay are incompletely understood.

While many proteins are involved in checkpoint maintenance, Aurora B Kinase serves as its master regulator [18, 19]. When the abscission checkpoint is satisfied, Aurora B activity at the midbody decreases and abscission occurs. In the presence of checkpoint-sustaining errors, Aurora B does not become inactivated at the midbody, and this cellular structure is instead stabilized. Aurora B maintains an active abscission checkpoint through phosphorylation of several proteins, including CHMP4C [22, 23]. Phosphorylated CHMP4C enforces abscission arrest by complexing with and sequestering the ATPase VPS4 needed to drive the membrane fission event of abscission [24–26]. VPS4 and CHMP4 proteins are subunits of the Endosomal Sorting Complexes Required for Transport (ESCRT) that complex with other ESCRT factors to facilitate membrane constriction and fission events, including cytokinetic abscission [27–29]. Additional abscission machinery, such as the early-acting ESCRT factor ALIX and the CHMP4C paralog CHMP4B, cooperate at the abscission site to complete cytokinesis [27, 28]. Upon abscission checkpoint resolution, CHMP4C releases VPS4, ALIX and CHMP4B are recruited to the abscission site, and membrane fission takes place [25].

Apart from DNA bridges, which provide a scaffold near the site of abscission where regulatory signaling can take place, conditions linked to abscission checkpoint activity, such as malformed nuclear pores or residual DNA damage, require a means by which to relay and/or amplify signals that keep the abscission checkpoint active. Under these circumstances, we discovered that tri-phosphorylated CHMP4C and a subset of Aurora B localize to cytoplasmic granules that we termed abscission checkpoint bodies (ACBs) [18, 30]. ALIX and CHMP4B are also found in ACBs, and the presence of ACBs is associated with delayed recruitment of ALIX to the midbody [30]. The persistence of ACBs was found to prolong time to abscission, supporting the hypothesis that ACBs contribute to an abscission delay. Surprisingly, ACBs were also found to contain the RNA splicing factor SRRM2.

SRRM2 localization to ACBs prompted the discovery that ACBs arise from mitotic interchromatin granules (MIGs), which are sites where nuclear speckle (NS) proteins concentrate at mitosis. NSs, a prevalent feature of nuclear architecture in interphase, comprise both long noncoding RNAs and hundreds of self-organizing proteins with roles critical to splicing, chromatin organization, and transcription [31]. At the onset of mitosis, NS-associated proteins disperse throughout the cytoplasm where, later in mitosis, they self-organize to form MIGs. MIGs are postulated to move splicing factors and small nuclear ribonucleoproteins (snRNPs) to daughter cells, thus ensuring their equal distribution [32, 33]. It is likely that ACBs arise from MIG precursors when abscission checkpoint signaling is sustained as ACBs are fewer in number and larger than MIGs. Although these two organelles have clear similarities in composition and properties, at least one compositional difference has been identified: ALIX is detected only in ACBs [30].

The functional significance of RNA splicing factors and NS/MIG components localizing to ACBs, an organelle with roles linked to the regulation of abscission, remains unknown. Here, we begin to decipher whether and how RNA splicing might participate in abscission and the abscission checkpoint. We show that select RNA splicing factors have altered localization when ACBs are present, and that the presence of ACBs is tightly correlated with decreased RNA splicing activity. Further, we show that inhibiting RNA splicing activity delays abscission, and we identify changes to transcription and splicing that occur at cytokinesis and during abscission checkpoint activity. Our findings demonstrate a reliance on RNA splicing for proper abscission timing, revealing how the coalescence of splicing factors with ESCRT proteins, and Aurora B kinase in ACBs contributes to abscission delay when the checkpoint is active.

## RESULTS

### ACBs are related to Nuclear Speckles and Mitotic Interchromatin Granules but have a distinct composition

To test whether SRRM2 is unique in its localization to ACBs, we probed for SRRM1 and SON and found that these splicing factors are also enriched in ACBs: (Figure 1A). Importantly, both ACBs and these resident proteins were detected in HeLa, DLD1, and U2OS cells and under distinct conditions that sustain abscission checkpoint activity: replication stress and reduced Nup153 levels [18, 21, 34, 35]. To investigate what distinguishes ACBs from MIGs, individual NS, MIG, and ACB components were tracked through the cell cycle. Whereas SRRM2 localized specifically to all three structures (NS, MIGs, and ACBs), other splicing factors, including SF2/ASF and SRSF7, were detected in MIGs but not ACBs (Figure 1B; Figure 1 Supplement 1A/B) [36]. SRRM2 levels (detected by the antibody mAbSC35 [37]) in MIGs increased under active checkpoint conditions (siNup153 treatment), whereas the SF2/ASF and SRSF7 signal intensities were equivalent in MIGs detected under siNup153 conditions compared to siControl. When ACBs form, SRRM2 signal in ACBs increased further compared to MIGs while SF2/ASF and SRSF7 intensities were lower in ACBs compared to their intensity in MIGs (Figure 1C/D; Figure 1 Supplement 1C-E). Consistent with SRRM2 cytoplasmic ACB localization, SRRM2 nuclear levels decreased as ACBs formed, while SF2/ASF and SRSF7 accumulated in nuclei (Figure 1E/F; Figure 1 Supplement 1F-H). These results underscore distinguishing features of MIGs and ACBs and reveal an altered repertoire of splicing factors in the nucleus when ACBs are present.

**Figure 1.**
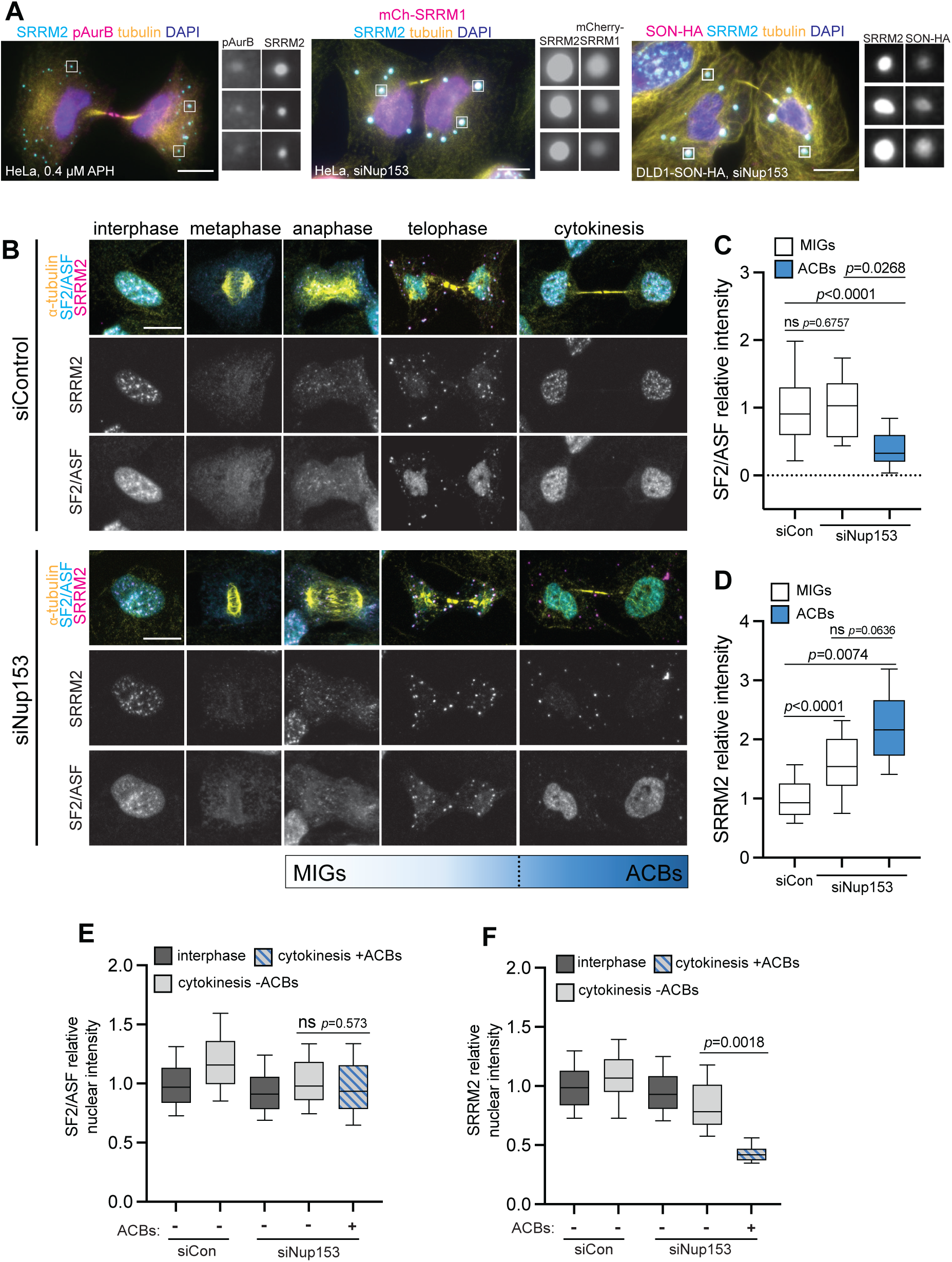
Splicing factors enrich differentially in mitotic interchromatin granules versus abscission checkpoint bodies. **A** Immunofluorescence images of cells under different treatments (0.4 µM Aphidicolin (APH), siNup153 as indicated) that sustain the abscission checkpoint and elicit ACBs. ACBs are tracked by detection of SRRM2 (mAbSC35), phospho-Aurora Kinase B, exogenously expressed mCherry-SRRM1, and endogenously tagged SON. Scale bars throughout all figures are 10 µm and ACB enlargements are 3 µm^2^. **B** Localization of SF2/ASF and SRRM2 (mAbSC35) in confocal Z-projections of pre-permeablized U2OS cells 48h following siControl (siCon)/siNup153 treatment. **C, D** Quantification of average SF2/ASF and SRRM2 (mAbSC35) intensity in MIGs and ACBs, normalized to the average siCon MIG intensity. Total n≥62 cells per experimental condition collected from 3 biological replicates. In all graphs with box and whiskers, center line represents mean and error bars represent 10-90% of distribution. **E, F** SF2/ASF and SRRM2 nuclear intensity normalized to average siCon interphase intensity. Total n≥166 cells per experimental condition collected from 3 biological replicates.

### SF3b1, a key RNA spliceosome component, is mislocalized when ACBs are present

To investigate further how RNA splicing may be altered under the abscission checkpoint, we turned our attention to the key RNA splicing factor SF3b1 (SF3b155). The N-terminus of SF3b1 scaffolds and binds the U2snRNP and is required for efficient splicing [38–40]. Knockdown or inhibition of SF3b1 changes RNA splicing by use of alternative branchpoints [41–44]. Therefore, changes to SF3b1 nuclear localization have the potential to alter RNA splicing.

SF3b1 localized both to NSs and throughout the nucleus during interphase but became diffuse throughout the cell in metaphase (Figure 2A). During telophase under unperturbed conditions, when SRRM2 targets to MIGs, SF3b1 localized to daughter nuclei. In contrast, under active checkpoint conditions, SF3b1 remained cytoplasmic during telophase. During cytokinesis, SF3b1 localized to the nucleus and NS under unperturbed conditions but was largely cytoplasmic when the abscission checkpoint was sustained by targeting Nup153 and Nup50 (Figure 2A, siNups). While SF3b1 was occasionally detected in ACBs, it did not consistently accumulate in these structures (Figure 2A). Compared to interphase cells, a significant decrease in nuclear enrichment of SF3b1 was detected in cytokinetic cells when ACBs were present, but not in siControl cytokinetic cells or in siNups cytokinetic cells that did not have ACBs (Figure 2B). To test whether reduced nuclear SF3b1 is linked to abscission checkpoint activity and the presence of ACBs, and not to a general nuclear pore defect, replication stress was used to sustain abscission checkpoint signaling. We again observed reduced nuclear SF3b1 in ACB-positive cells (Figure 2 Supplement 1A-C). Importantly, neither Nup153/50 depletion nor replication stress reduced SF3b1 nuclear localization at interphase (Figure 2B and Figure 2 Supplement 1A-C).

**Figure 2.**
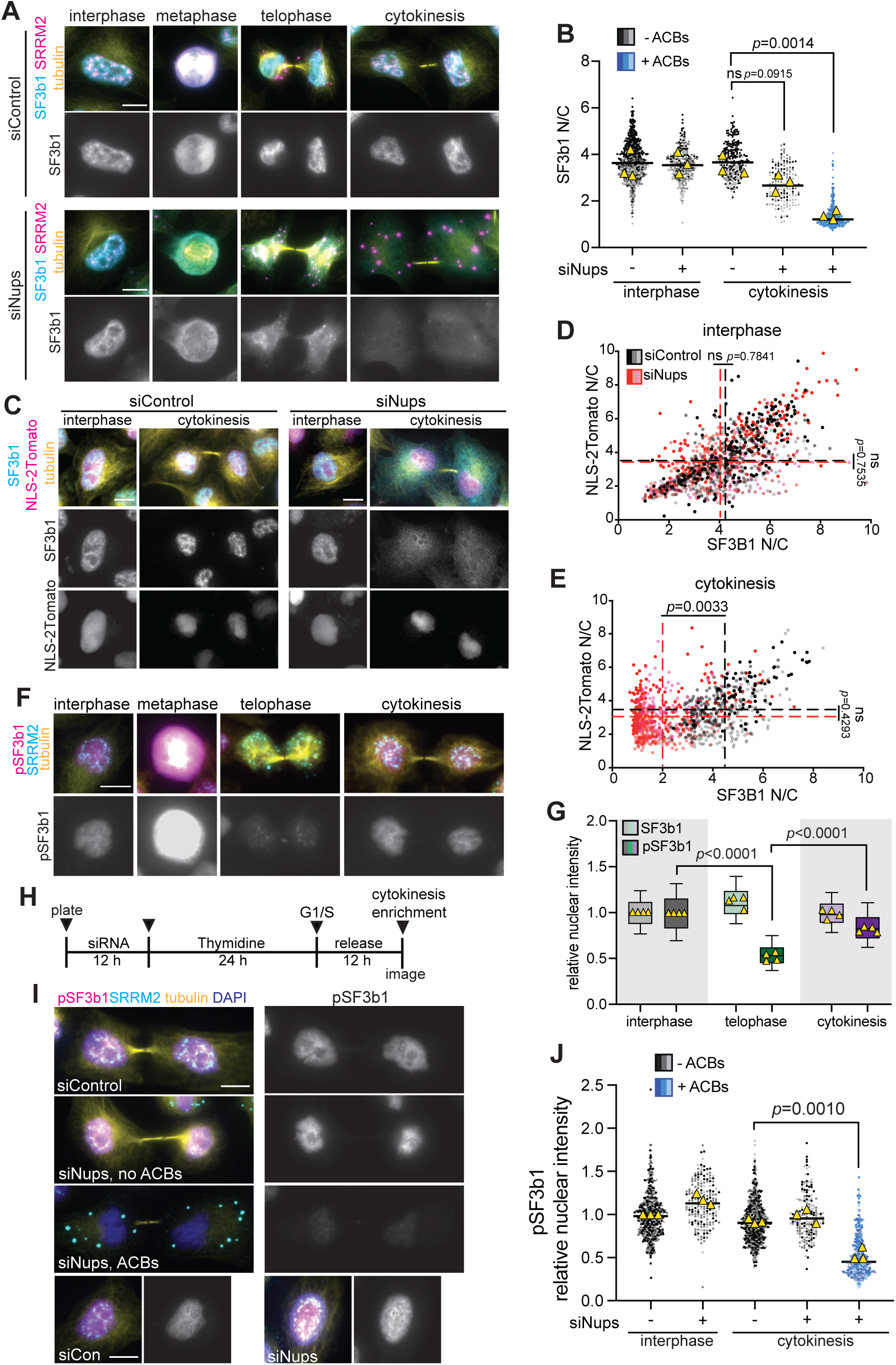
SF3b1 localization and activity are altered when ACBs are present. **A** SF3b1 and SRRM2 (mAbSC35) in immunofluorescence images of HeLa cells through the cell cycle with siCon/siNup50 and siNup153 (siNups) treatment. **B** Quantification of nuclear-to-cytoplasmic (N/C) intensity of SF3b1 in interphase and cytokinetic cells treated with siCon/siNups. Total n≥286 cells per experimental condition collected from 3 biological replicates. Means from each biological replicate are shown in triangles and black lines represent the mean of all replicates in all figures. **C** Immunofluorescence images of HeLa cells expressing an exogenous NLS peptide fused to two tandem Tomato fluorescent proteins and stained for SF3b1. **D, E** Quantification of N/C intensities of NLS-2Tomato and SF3b1 in siCon/siNups treated interphase (D) and cytokinetic (E) cells. Total n≥389 cells per experimental condition collected from 3 biological replicates. Dashed black lines indicate siCon treated NLS-2Tomato and SF3b1 N/C averages and red dashed lines indicate siNups NLS-2Tomato and SF3b1 N/C averages. **F** Immunofluorescence images of pSF3b1(Thr313) and SRRM2 in HeLa cells at various stages of the cell cycle. **G** Quantification of nuclear intensity of SF3b1 and pSF3b1 in interphase, telophase and cytokinetic cells normalized to interphase average nuclear intensity. Total n≥170 cells per experimental condition collected from 4 biological replicates. **H** Treatment time course of HeLa cells under thymidine synchronization for cytokinesis enrichment and treatment of siCon/siNups (siRNA, 36 h). **I** Immunofluorescence images of pSF3b1 and SRRM2 in HeLa cells at cytokinesis (6 upper panels) and interphase (4 lower panels) treated as in (H). **J** Quantification of pSF3b1 nuclear intensity of HeLa cells (treated as in H). pSF3b1 nuclear intensity was normalized to average siCon interphase nuclear intensity in each individual experiment. Total n≥198 cells per experimental condition collected from 3 biological replicates.

To test whether SF3b1 mislocalization was due to a global impairment of post-mitotic nuclear import when Nup153 and Nup50 were depleted, we concomitantly tracked a fluorescently tagged nuclear import reporter (NLS-2Tomato) (Figure 2C). Both NLS-2Tomato and SF3b1 localized predominantly to the nucleus under siControl and siNups conditions at interphase, with no significant difference (Figure 2C/D). At cytokinesis, when SF3b1 remained cytoplasmic in siNups-treated cells, nuclear accumulation of the NLS reporter remained equivalent, indicating that nuclear import machinery operates in abscission checkpoint active cells (Figure 2C/E). We also tested whether the extent of cytoplasmic accumulation of SF3b1 might correspond to levels of Nup153/Nup50 depletion but found the same range of knockdown efficiency in cell populations with and without SF3b1 cytoplasmic persistence (Figure 2 Supplement 1F-I). Hence, SF3b1 cytoplasmic localization corresponded most tightly to the presence of ACBs (Figure 2B and Figure 2 Supplement 1C), rather than reflecting a general defect in nuclear import or variations in Nup153/50 depletion.

### SF3b1 does not associate with catalytically active spliceosomes when ACBs are present

To understand the biological impact of altered SF3b1 distribution, we first sought to determine whether splicing is normally underway as cells progress from telophase to the completion of cytokinesis. We monitored phosphorylation of SF3b1 at Thr313 (pSF3b1), known to be targeted by CDK11 concomitantly with spliceosome catalytic activity [45–47]. In untreated, asynchronous cells, pSF3b1 nuclear intensity was detected in interphase cells. pSF3b1 levels increased further during prometaphase and metaphase, consistent with this site also being a mitotic target of CDK1 [48]. Despite this strong peak of mitotic phosphorylation, levels of pSF3b1 markedly decreased at telophase, before returning to interphase-levels at cytokinesis in cells still connected by a midbody structure. The latter result indicates that spliceosome activity resumes before cells undergo abscission (Figure 2F/G). The decrease in pSF3b1 intensity in telophase was not due to lower overall levels of SF3b1, as total SF3b1 nuclear intensity was unchanged at telophase compared to interphase or cytokinesis (Figure 2A, Figure 2F/G).

To assess the status of the residual SF3b1 nuclear population that remains when the abscission checkpoint is sustained, cells subjected to siControl or siNups conditions were incubated in thymidine to arrest the cell cycle in G1/S, followed by a 12-hour release to enrich for cytokinetic cells (Figure 2H) [30]. Under these conditions, pSF3b1 intensity was reduced specifically in cells positive for ACBs (Figure 2I/J). Similarly, ACB-positive cells that arose following replication stress also had reduced nuclear pSF3b1 levels (Figure 2 Supplement 1A/D/E). In cells treated with siNups, cytokinetic cells with high and low levels of nuclear pSF3b1 shared similar levels of Nup50 and Nup153 depletion (Figure 2 Supplement 1J-M). Together these results demonstrate that the reduced levels of nuclear SF3b1 and pSF3b1 track together. Thus, any splicing taking place in ACB-positive cells occurs with reduced levels of the cofactor SF3b1, which is likely to result in an altered RNA landscape [41–44].

### RNA splicing inhibition delays abscission

To test whether RNA splicing contributes to abscission progression and whether altered SF3b1 activity mediates abscission delay, HeLa cells were treated with inhibitors that specifically target activity of the SF3B complex, Pladienolide B (PB) [49] and Spliceostatin A (SSA) [50], or a DMSO vehicle control. HeLa cells stably expressing GFP-tubulin were live-imaged under asynchronous growth conditions, and cells entering telophase 2-3 hours following the addition of inhibitor were analyzed (Figure 3A). Progression from prophase, identified by the first frame in which microtubules were seen to nucleate around centrosomes (Figure 3B, blue arrows), to telophase, identified by the first frame in which a midbody structure can be seen (Figure 3B, white arrows), was measured. Under all treatment conditions, cells had similar kinetics of mitotic transit (DMSO: t_50_=45 min; PB: t_50_=45 min; SSA: t_50_=44 minutes, Figure 3C). In contrast, progression through abscission, the time from midbody formation to its disassembly (Figure 3B, green arrows), was significantly delayed by the SF3b1 inhibitors (DMSO: t_50_=74 min; PB: t_50_=92 min, p=0.0289; SSA: t_50_=94 min, p=0.0435 Figure 3D). Thus, abscission timing is delayed when mRNA splicing is disrupted, recapitulating events that occur when ACBs mediate checkpoint activity.

**Figure 3.**
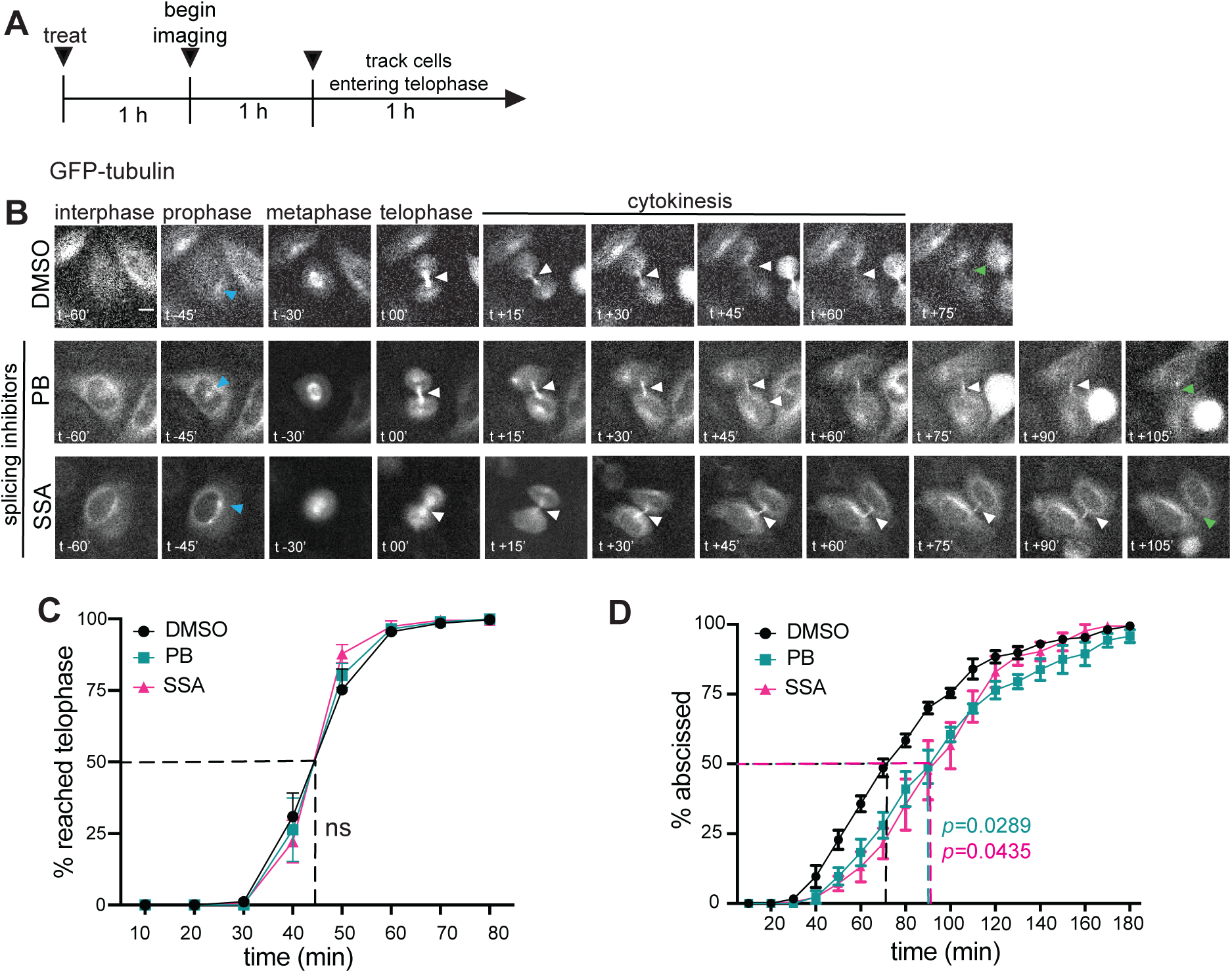
RNA splicing inhibition delays abscission but not mitosis. **A** Time course of DMSO (n=204), Pladienolide B (PB, 10 µM) (n=141), or Spliceostatin A (SSA, 0.2 µM) (n=113) treatment and live imaging of asynchronous HeLa cells from four biological replicates. **B** Live imaging of GFP-tubulin-expressing HeLa cells. Images were independently adjusted for brightness and contrast. Prophase was identified by microtubule clustering (blue arrows), telophase and cytokinesis were identified by the formation of a midbody structure (white arrow), and abscission was identified by a separated midbody or the frame in which the midbody is no longer visible (green arrows). **C, D** Quantification of time from microtubule clustering (prophase, blue arrows) to midbody formation (telophase, white arrows) (DMSO: t_50_=44.6 min, PB: t_50_=44.9 min, SSA: t_50_=44 minutes) (C), and time from midbody formation (telophase, white arrows) to loss of midbody (abscission, green arrows) (DMSO: t_50_=74 min, PB: t_50_=93 min, SSA: t_50_=94 min) (D).

### The abscission checkpoint is associated with changes in gene expression

The alterations in RNA splicing factor localization and activity under the abscission checkpoint prompted us to investigate whether overall RNA biogenesis is disrupted when the checkpoint is active. To evaluate the RNA repertoire of ACB-containing cells, we performed RNA-sequencing on cell populations following a mitotic shake-off method to enrich for cells at cytokinesis under active and inactive (siNups/siCon) checkpoint conditions, and their interphase-stage counterparts (Figure 4A/B, Figure 4 Supplement 1A). Principal component analysis revealed that each of the four different combinations of cell cycle stage and Nups depletion status exhibited a unique gene expression pattern (Figure 4 Supplement 1B). Analysis of differential gene expression examining the interaction between Nups depletion status and cell cycle stage revealed 627 genes with >1.5-fold changes (Figure 4C). From this gene set, we were able to identify genes that were uniquely influenced by Nup50/153 depletion during cytokinesis. RNAs specifically downregulated under the abscission checkpoint (siNups, cytokinesis) were enriched in biological processes involved in transcription (Figure 4C, clusters 1 and 2; Figure 4D). This was also true of gene sets that were downregulated at cytokinesis regardless of Nup50/153 status (Figure 4 Supplement 2A/B). A substantial portion (246 transcripts) of the 627 genes that were differentially expressed in response to the interaction of Nup153/50 status and cell cycle stage were found to be elevated under normal cytokinetic conditions but failed to do so under checkpoint-sustaining siNups treatment (Figure 4C, cluster 3). While there was no clear enrichment of genes that share biological functions in this cluster, a notable feature was the enrichment of lncRNAs, which comprised 55% of this RNA population (Figure 4E). lncRNAs were also significantly upregulated more generally during cytokinesis regardless of Nups status (Figure 4 Supplement 2A/C). Genes differentially expressed in response to siRNA treatment independently of cell cycle stage were evaluated in Figure 4 Supplement 2 D-F. Overall, our data reveal that the cellular RNA profile changes during cytokinesis, and that this profile is altered further when the abscission checkpoint is sustained by Nup153/50 depletion.

**Figure 4.**
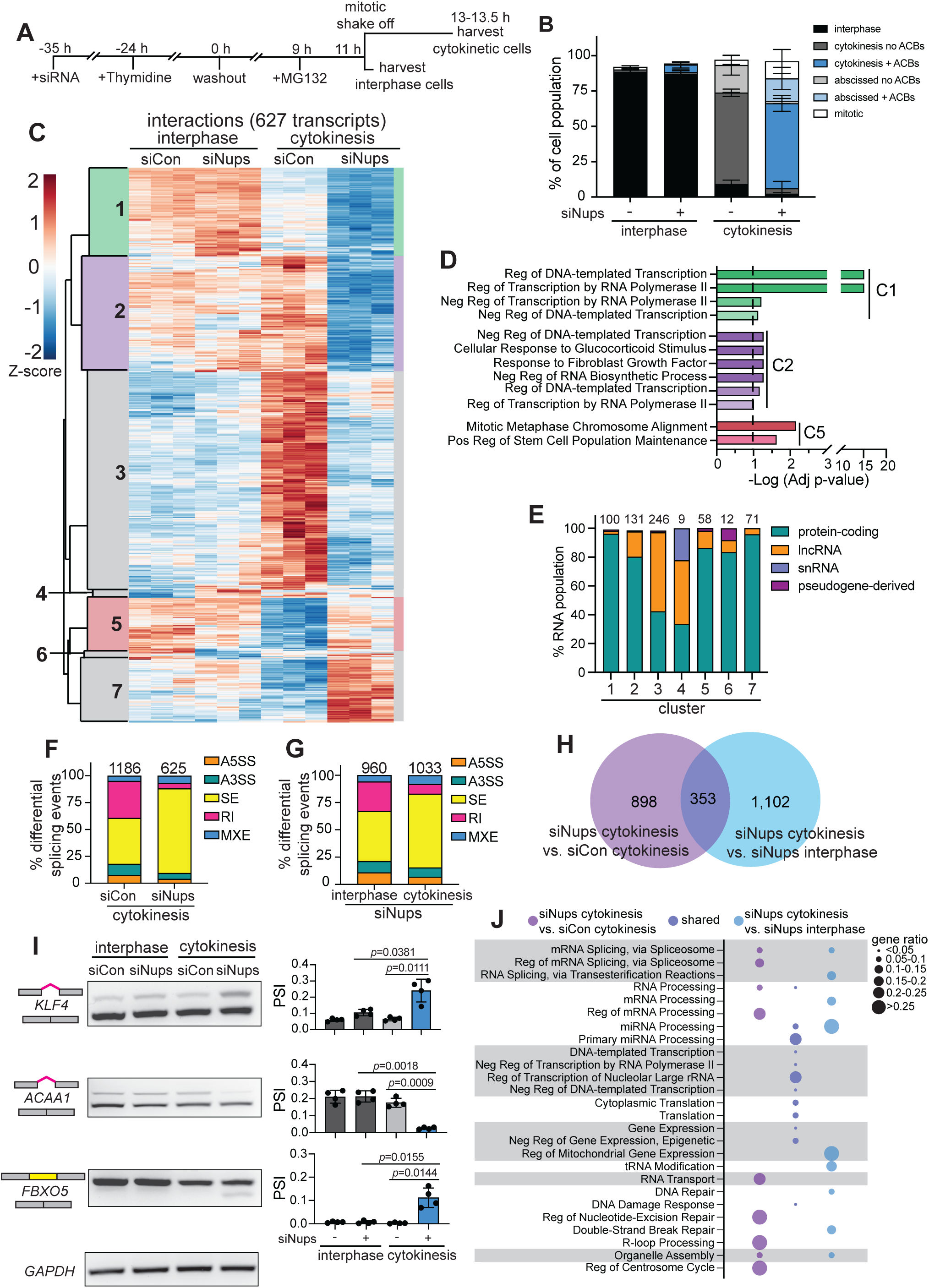
Gene expression and RNA splicing are altered when the abscission checkpoint is active. **A** Treatment time course to collect HeLa cells expressing GFP-tubulin and mCherry-SRRM1 at interphase and cytokinesis treated with either siCon or siNups. **B** Quantification of fluorescent images of interphase and cytokinetic cell populations (treated as in A). Total n≥720 cells per experimental condition collected from 3 biological replicates. **C** Clustered heatmap of 627 differentially expressed genes that had significant interactions between cell cycle state (interphase/cytokinesis) and siRNA treatment (siCon/siNups) (|fold-change|≥1.5, p_adj_<0.05). Z-scores reflect standard deviations above (positive, red) or below (negative, blue) average regularized log-transformed read-counts across samples. **D** Biological processes gene ontologies significantly enriched (p_adj_<0.1) in cluster 1, 2, and 5 transcripts (C). Gene ontology analysis of Clusters 3 and 7 did not reveal any statistically significant enrichments. Clusters 4 and 6 contained only 9 and 12 transcripts respectively, so gene ontology analysis was not performed. Dashed line represents p_adj_=0.1. **E** The proportion of RNA biotypes identified in clusters 1-7 (C). Biotypes with <2% representation were omitted. Numerical values above bars represent the number of transcripts in each cluster. **F** Proportion of significant alternative splicing events with |proportion spliced in (PSI)|≥0.1 between siNups cytokinesis and siCon cytokinesis cells (false discovery rate (FDR)<0.05). Numerical values represent each differential splicing event from the comparison that occurs in either condition. **G** Proportion of significant (FDR<0.05) alternative splicing events with |PSI|≥0.1 between siNups cytokinesis and siNups interphase cells, as in F. **H** Significant alternative splicing events (FDR<0.05, |PSI≥0.1) that are unique to the siNups cytokinesis/siCon cytokinesis comparison (purple, 1,251 events), unique to the siNups cytokinesis/siNups interphase comparison (light blue, 1,455 events), or shared between these comparisons (dark purple, 353 events). **I** RT-PCR (left) and densitometric analyses (right) of significantly alternatively spliced transcripts (KLF4, retained intron 2; ACAA1, retained intron 4; FBXO5, skipped exon 3) identified in H (shared events, dark purple). GAPDH was used as an unspliced loading control. **J** Biological processes gene ontologies of interest significantly enriched (p_adj_<0.1) in alternatively spliced targets identified from comparisons show in H (siNups cytokinesis v. siCon cytokinesis, siNups cytokinesis vs. siCon interphase, shared). GOs are grouped by biological process similarity, and gene ratios are depicted by circle size.

### The abscission checkpoint is associated with changes in RNA splicing

We next analyzed RNA splicing events corresponding to abscission checkpoint activity. Because of the decrease of nuclear pSF3b1 and the observed cytoplasmic enrichment of SF3b1, SON, SRRM2, and SRRM1 when ACBs arise, and splicing changes that occur when SF3b1 is altered [41–44], we predicted that RNA splicing would be affected during the abscission checkpoint. Indeed, we observed increased proportions of significant differential skipped exon (SE) splicing events and decreased proportions of significant retained intron (RI) events when siNups cytokinetic cells were compared to both siControl cytokinetic cells and siNups interphase cells (Figure 4F/G; see Figure 4 Supplement 1C/D for additional comparisons). At the same time, we observed a reduced overall number of significant differential splicing events in the siNups cytokinetic cells (625 events) vs. siControl cytokinetic cells (1,186 events), potentially reflecting generally reduced splicing activity (Figure 4F).

To home in on splicing changes specific to the abscission checkpoint, we eliminated events that could be attributed to Nup153/50 depletion (i.e. present in the siNups interphase vs. siControl interphase comparison) and to cytokinesis-specific events (i.e. present in the siControl cytokinesis vs. siControl interphase comparison) (Figure 4 Supplement 1E/F). Of the remaining 1,455 alternative splicing events that occurred in the siNups cytokinesis vs. siNups interphase comparison, and the 1,251 events that occurred in the siNups cytokinesis vs. siControl cytokinesis comparison, 353 events were shared (Figure 4H). Three mRNAs from this group were tracked individually by RT-PCR to validate the bioinformatic analyses. We confirmed increased intron 2 retention of KLF4 (KLF4, top band), decreased intron 4 retention of ACAA1 (ACAA1, top band), and skipping of exon 3 of FBXO5 (FBXO5, bottom band) in Nup153/50-depleted cytokinetic cells (Figure 4I). Thus, an altered program of splicing ensues when the abscission checkpoint is active.

Gene Ontology (GO) enrichment analysis of transcripts affected by the 353 alternative splicing events specific to the abscission checkpoint revealed significant enrichment in biological processes related to gene expression, transcription, translation, and DNA damage response among others (Figure 4J). These biological processes, in addition to several related to splicing, also emerged when looking more broadly at alternative splicing within comparisons that enriched for abscission checkpoint conditions (i.e. siNups cytokinetic cells vs. siNups interphase cells and siNups cytokinetic vs. siControl cytokinetic cells, Figure 4J), as well as during cytokinesis more generally (i. e. siControl cytokinetic vs. siControl interphase cells, Figure 4 Supplement 1G). Therefore, trends in gene expression and splicing that are present as cells undergo cytokinesis under unperturbed conditions are exacerbated under the abscission checkpoint. More generally, the prevalence of biological processes related to RNA processing among genes that are alternatively spliced when abscission checkpoint activity is sustained indicates that the abscission checkpoint alters RNA biogenesis through both decreased gene expression and alternative splicing.

## DISCUSSION

Our previous discovery of SRRM2 in ACBs [30] suggested a novel relationship between RNA splicing and the abscission checkpoint, yet the functional significance of SRRM2 sequestration in ACBs remained unclear. Here, we reveal that RNA splicing is required for timely abscission progression, and that splicing status is modulated when ACBs form owing to abscission checkpoint activity. We also identified SRRM1 and SON as ACB residents. Given that SRRM2 and SON provide structural scaffolding for NS assembly [37, 51], these proteins likely provide similar structural support to ACBs. In contrast, SF2/ASF and SRSF7 were not enriched in ACBs, despite their documented localization to MIGs (Figure 1B/C, Figure 1 Supplement 1) [36]. Therefore, although ACBs share properties and components with NS and MIGs, they are all distinct organelles. During telophase, some proteins leave MIGs and return to the nucleus, whereas other proteins such as ALIX are targeted specifically to ACBs [30]. Correspondingly, MIGs and ACBs apparently have specialized functions, as MIGs are hypothesized to be sites of RNA splicing factor modification and storage at mitosis [32, 33], whereas we propose that an important aspect of ACB function is to delay abscission by sequestering splicing factors.

Cytoplasmic localization of SF3b1 is not dictated by the extent of Nup153 or Nup50 depletion per se, but rather is tightly associated with the presence of ACBs (Figure 2A/B, Figure 2 Supplement 1A-C). Cytoplasmic SF3b1 localization does not appear to reflect impaired core functions of the nuclear pore or loss of nuclear integrity because an import reporter bearing a canonical NLS remains robustly imported under these same conditions (Figure 2C-E). Conceivably, however, Nup153 and Nup50 could be selectively important for the efficiency of SF3b1 import in post-mitotic cells, with abscission checkpoint activity and ACB biogenesis contributing cooperative signals that synergistically impair import of SF3b1 and other cargo such as SRRM2 and SON. Consistent with this idea, we observed a trend of delayed nuclear accumulation of SF3b1 in siNups-treated midbody-stage cells lacking ACBs (Figure 2B). Alternatively, the trend in delayed SF3b1 import in ACB-negative cells could reflect cells that have recently satisfied the abscission checkpoint and are progressing to abscission.

In support of the hypothesis that altered RNA splicing can contribute to delayed abscission under an active checkpoint, we found that pharmacological inhibition of RNA splicing in normally cycling cells delays abscission, and that splicing is therefore required for efficient abscission (Figure 3B/D). However, the extent of delay (20-25 min) was not as robust as when the checkpoint is sustained through siNup153 depletion (3.25-4 h) [18]. The weaker phenotype observed when splicing is altered in isolation may be due to the absence of cooperative signals that also lead to AuroraB kinase signaling and CHMP4C regulation [18, 19, 22, 23]. We therefore speculate that altered RNA splicing works in combination with other regulators of the abscission checkpoint to delay abscission.

Transcriptomic analyses revealed specific downregulation of genes enriched in multiple biological processes related to transcription specifically in cell populations containing primarily cytokinetic cells with active abscission checkpoints (siNups cytokinesis) (Figure 4C/D). These data indicate that both RNA splicing activity and RNA biogenesis more generally may promote abscission. Additionally, we identified a set of genes that are upregulated as cells progress through cytokinesis without checkpoint activity (siCon cytokinesis). Interestingly, many of the transcripts in this cluster were classified as lncRNAs (Figure 4E). This finding reveals an underappreciated set of genes with roles that may be important to mitotic exit and progression through abscission.

Recent publications have illuminated the process of midbody-localized translation and its importance for efficient cytokinetic abscission, further highlighting the role of RNA biogenesis during cytokinesis [52, 53]. We did not identify consistent correspondence when comparing differentially expressed genes in our study to midbody-enriched transcripts. However, 6.5-18% of transcripts alternatively spliced during the abscission checkpoint (Figure 4H) also enrich at the midbody when compared to published datasets [52, 53]. This correspondence could point to a mechanism by which alternative splicing affects proteins produced by midbody-localized translation.

While alternatively spliced transcripts can be unproductive and undergo degradation by nonsense-mediated mRNA decay [54], we did not find correspondence between RNA levels and alternatively spliced RNAs. Instead, our findings indicate that an altered protein repertoire works to pause the cell cycle when the abscission checkpoint is active, building upon a limited number of studies that have characterized alternatively spliced protein isoforms as abscission regulators [55, 56]. Here, the data obtained open new lines of investigation to map how specific splice products contribute to abscission and its regulation.

## METHODS

### Cell culture

HeLa and U2OS cells were cultured and maintained at 37°C and 5% CO_2_ in DMEM supplemented with 10% FBS. DLD1 cells were cultured and maintained at 37°C and 5% CO_2_ in DMEM supplemented with 10% FBS and GlutaMax (Thermo Fisher Scientific, Waltham, MA). At the outset of these studies, HeLa, U2OS, and DLD1 cells were tested and found negative for mycoplasma using a PCR mycoplasma detection kit (ABM, Bellingham, WA). Transduced cell lines expressing exogenous constructs were validated by genomic testing. Unless otherwise labeled in individual panels or figure legends, all experiments used HeLa cells.

### siRNA transfections

Cells were transfected with siRNA for 48 h, as indicated in figure legends, using Lipofectamine RNAiMax (Thermo Fisher Scientific, Waltham, MA) according to the manufacturer’s instructions. Media was exchanged 8-24 h after transfection, and cells were incubated between 18 and 48 h before harvesting.

### Immunofluorescence

Cells were seeded on fibronectin-coated (10 µg/mL) glass coverslips or directly onto glass coverslips and washed once with 1X PBS before fixation with one of three techniques: 1) 4% paraformaldehyde (PFA) 23 °C for 15 minutes and rinse with PBS, followed by permeabilization with 0.5% Triton-X in PBS at 23 °C for 5 minutes; 2) rinse with 50 mM PIPES, 25 mM HEPES pH 7.0, 10 mM EGTA, 4 mM MgSO_4_, with PMSF [1 mM], aprotinin, leupeptin added freshly (PHEM) buffer at 23 °C, pre-permeabilization with 0.5% Triton-X in PHEM buffer for 1 min at 23 °C, fixation with 2–4% PFA at 23 °C for 15 min, rinse with PBS, and incubation with 0.5% Triton-X in PBS for 5 min at 23 °C; or 3) fixed with cold MeOH at −20 °C for 8 minutes and rinsed with PBS. Following fixation, coverslips were coated in blocking solution (3% FBS, 0.1% Triton-X in PBS) for 30 min at 23 °C. Primary antibodies were applied at the dilutions noted (key resources table) for at least 1 h, 23 °C in blocking solution. After 1 wash with PBS, secondary antibodies (Thermo Fisher Scientific) were applied for 45-60 min, and cells were washed in PBS. Coverslips were mounted with ProLong Glass Antifade Reagent with NucBlue (Thermo Fisher Scientific) on a microscope slide. Coverslips were immediately sealed with clear nail polish and cured at 23 °C in the dark overnight, before moving to short-term storage at 4 °C or long-term storage at −20 °C.

### Nup depletion

HeLa cells were seeded onto fibronectin-coated (10 µg/mL) glass coverslips or directly onto glass coverslips at a density of 55-75,000 cells/mL (Figure 1A, Figure 1Supplement 1A/B, Figure 2A/C, Figure 2 Supplement 1F/G) and U2OS cells were seeded at a density of 100,000 cells/mL (Figure 1B). Seeded cells were treated with 10 nM non-targeting siRNA (siCon) or siRNAs targeting Nup153 (Figure 1B, Figure 1 Supplement 1A/B) or Nup153 and Nup50 (Figure 2A/C, Figure 2 Supplement 1F/G) (siRNA sequences previously used in [30]). 24 h post-transfection the media was replaced, and 48 h post-transfection cells were fixed with PHEM followed by 4% PFA (Figure 1B, Figure 1 Supplement 1A/B) or 4% PFA (Figure 1A, Figure 2A/C, Figure 2 Supplement 1F/G) and labeled with antibodies as described above.

In synchronized experiments, HeLa cells were plated at 70,000 cells/mL 48 h prior to fixation and treated with siCon or siNup50 and siNup153 in combination (10 nM each). 36 h prior to fixation, 2 mM thymidine was added to cells in fresh media. 12 h prior to fixation, cells were washed with PBS thrice and media was replenished. Cells were fixed with 4% PFA and antibody labeled as described above (Figure 2H, Figure 2 Supplement 1J/K).

### Replication stress assays

HeLa cells were seeded at 100,000 cells/mL onto fibronectin-coated (10 µg/mL) glass coverslips and synchronized with 2 mM thymidine. 24 h post-seeding cells were washed three times with PBS and replenished with fresh media containing 0.4 µM Aphidicolin (APH) or in 0.04% DMSO. 13 h after the addition of APH, cells were fixed with 4% PFA and immunofluorescence labeling was performed as described above (Figure 2 Supplement 1B/D).

### Untreated and asynchronous HeLa cell experiments

HeLa cells were seeded at 100,000 cells/mL onto fibronectin-coated (10 µg/mL) glass coverslips 24 h prior to fixation. Cells were fixed with 4% PFA and immunofluorescence labeling was performed as described (Figure 2F).

### Live cell imaging

4-well chamber coverslips (Lab-Tek, Wilmington, NC) were coated with fibronectin (10 µg/mL) for 10 minutes. HeLa cells stably and constitutively expressing GFP-tubulin were seeded onto the chamber coverslips at a concentration of 100K cells/mL 24 h prior to the start of live imaging. 8 h prior to imaging, media was replaced with live imaging-compatible media (phenol-free DMEM/F-12 buffered with HEPES, or phenol-free Lebovitz’s L-15 media). 1 h prior to imaging, cells were treated with 0.01-1% DMSO, 10 µM PB, or 0.2 µM SSA. Imaging was carried out on a Nikon Ti-E widefield inverted microscope (Nikon 20x N.A. dry objective lens) equipped with Perfect Focus system and housed in a 37°C chamber (OKOLAB, Ambridge, PA) with 5% CO_2_. 10 fields of view per treatment condition were selected at various x and y coordinates, and images were acquired using a high-sensitivity Andor Zyla CMOS camera (Andor, Manchester, CT) controlled by NIS-Elements software. The treated cells were imaged for 12 h at intervals of 5 minutes. Times from mitosis to telophase, and telophase to abscission were measured in cells entering telophase 2-3 hours post-inhibitor treatment (Figure 3A/B).

### Image acquisition

Images were acquired using three different microscopes:

1. Leica SP8 Confocal 63× 1.4 oil HC PL APO objective with adjustable white-light laser to control for bleed-through (Figure 1B, Figure 1 Supplement 1A/B). Images were acquired as z-stacks and each individual slice was deconvolved using the Hyvolution and Lightening modes within the Leica App Suite X Software. Presented images are maximum z-projections of the deconvolved slices.
2. Zeiss Axioskop 2 Widefield 63× 1.4 oil DIC Plan Apochromat objective (Figure 1A, Figure 2A/C/F/I, Figure 2 Supplement 1B/D/F/G/J/K, Figure 4 Supplement 1A). Images were acquired as single plane widefield images.
3. Nikon Ti-E widefield inverted microscope (Nikon 20x N.A. 0.8 objective lens) and Andor Zyla CMOS camera (Andor, Manchester, CT) (Figure 3B). Live cell images were acquired as single plane widefield images.

### Cell and image scoring

Midbody-stage cells were identified by α-tubulin staining and were always counted as one cell. Cell populations were counted and sorted into interphase, cytokinesis, telophase, mitotic, recently abscised, multinucleate, and failed bridge (counted as one cell). Cells were counted from single-plane images acquired using the Zeiss Axioskop 2 Widefield 63× 1.4 oil DIC Plan Apochromat objective.

Midbodies were designated as ‘early’ (telophase) or ‘late’ (cytokinesis) using α-tubulin staining, and this was done prior to scoring for the examined phenotype. Midbodies were classified as early based on midbody width, level of midbody-pinching, nuclear area, and flatness of cells. Telophase-stage cells were identified as having condensed DNA and a thick tubulin-dense bridge (early midbody); whereas cytokinetic cells were identified as having decondensed nuclear DNA, a larger cytoplasmic area surrounding the daughter nuclei, and a thin tubulin bridge (late midbody). Cytokinetic cells were scored as having ACBs if they had at least one ACB. However, most cytokinetic cells with ACBs had 10–40 ACBs.

SF2/ASF, SRSF7, and SRRM2 intensity in MIGs and ACBs was measured using Zen software (Zeiss). Telophase and cytokinetic cells were identified by eye and measured individually. MIGs and ACBs were automatically gated on SRRM2 extra-nuclear signals in telophase and cytokinetic cells, and SRRM2, SF2/ASF, and SRSF7 average intensities within the SRRM2-positive foci were measured automatically. Three random cytoplasmic regions were selected in each cell to identify background levels of SRRM2, SF2/ASF, and SRSF7 in the areas surrounding MIGs and ACBs. These background measurements were averaged and subtracted from the MIG and ACB intensities in each cell to show overall enrichment in MIGs and ACBs versus the surrounding cytoplasm (Figure 1C/D, Figure 1 Supplement 1C-E).

SF3b1 and NLS-2Tomato nuclear:cytoplasmic intensities (N/C) were measured using Zen software (Zeiss). The nucleus was gated on the DAPI channel and average nuclear intensities of SF3b1 and NLS-2Tomato were reported. SF3b1 and NLS-2Tomato cytoplasmic intensities were measured from regions surrounding the nucleus (Figure 2B/D/E, Figure 2 Supplement 1C).

Nuclear intensities of SF2/ASF, SRSF7, SRRM2, Nup153, Nup50, SF3b1, and pSF3b1 were measured using Zen software (Zeiss). The nuclear region was gated on DAPI and average Nup153, Nup50, SF3b1, or pSF3b1 intensity was measured in this defined region (Figure 1E/F, Figure 1 Supplement 1F-H, Figure 2G/J, Figure 2 Supplement 1E/H/I/L/M).

For semi-quantitative RT-PCR image analysis, densitometric ratios were calculated using Fiji software (NIH). Gel bands were selected and plotted using the “Analyze Gels” feature in the Fiji analysis software, and band intensities were measured. The proportion of intron retained:intron spliced, or exon skipped:exon retained was then calculated (Figure 4I).

### Mitotic shake off for cytokinesis enrichment

HeLa cells constitutively expressing GFP-tubulin and mCherry-SRRM1 were plated at a density of 120,000 cells/mL in 10 cm dishes, containing two fibronectin-coated (10 µg/mL) glass coverslips. Cells were treated with siControl or siNups siRNAs for 11 hrs. Media was then replaced with media containing 2 mM thymidine. 24 h post-thymidine addition, cells were washed thrice with PBS and media was replaced. 9 h post-thymidine washout media was gently aspirated and replaced with media containing 5 µM MG132. 2 h post-MG132 addition, media was gently pipetted up and down on the plate 8 times and collected to isolate mitotic cells. The cells remaining on the plate were washed thrice with PBS to further remove mitotic cells, coverslips were fixed with 2% PFA for microscopy and mounted to slides for microscopic analysis and the remaining interphase cells were lysed for RNA extraction. The collected mitotic cells were pelleted and washed with 13 mL PBS at 4° C twice before resuspension in 2.5 mL warmed media. 300 µL of resuspended mitotic cells were plated onto fibronectin-coated (10 µg/mL) glass coverslips for microscopic analysis and the remaining 2.2 mL were plated into a single well of a 6-well plate for RNA extraction. Mitotic cells were released for 2.25-2.5 h, when most cells had reached cytokinesis by visual inspection. Cells were then gently washed once with PBS and either fixed with 2% PFA for and mounted to slides microscopic analysis or lysed for RNA extraction (Figure 4 A/B, Figure 4 Supplement 1A).

### RNA extraction, cDNA synthesis, and PCR

Total cellular RNA was isolated from cells under four conditions as described above (siControl interphase, siNups interphase, siControl cytokinesis, and siNups cytokinesis) using the Monarch Total RNA Miniprep Kit (New England Biolabs, Ipswich, MA). The included on-column DNase treatment was performed on all collected samples following manufacturer’s instructions. 10 ng input RNA was used, and cDNA synthesis and PCR were performed in combination using the OneTaq One-Step RT-PCR Kit (New England Biolabs). Following a 30-minute reverse transcription step at 48 °C, PCR reactions were performed under the following conditions: initial denaturation for 1 minute at 94 °C, 30 cycles with 15 seconds at 94 °C, 30 seconds at 55 °C, and 1 minute at 68 °C, followed by a final extension of 5 minutes at 68 °C. PCR products were then resolved on 2% agarose gels. The primer sequences used can be found in the table below (Figure 4I).

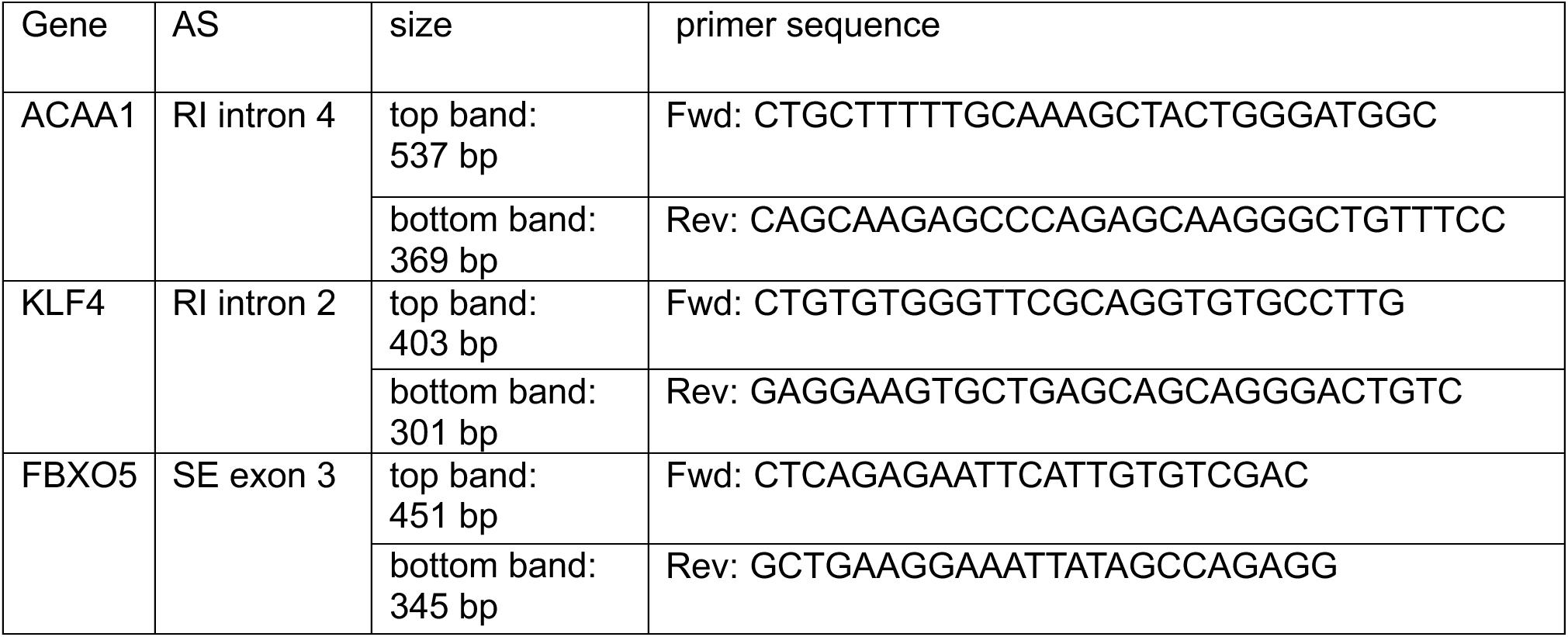

### RNA sequencing, differential gene expression and splicing analysis

Total RNA was isolated as described above. RNA quality control, cDNA library preparation, and sequencing were performed by the University of Utah High Throughput Genomics Shared Resource. RNA quality control was performed using the Agilent RNA ScreenTape Assay (Agilent Technologies, Santa Clara, CA). mRNA was isolated and a cDNA library was created using the NEB Ultra II directional RNA Library Prep with Poly(A) mRNA Isolation (New England Biolabs, Ipswich, MA). RNA alignments and counts were performed by the University of Utah Cancer Bioinformatics Shared Resource. Prior to alignment, adapters were trimmed using cutadapt (v. 3.4) [57]. Alignments were done using STAR (v. 2.7.9a) [58], and statistics were collected with Picard CollectRnaSeqMetrics (v. 2.26.2) (Broad Institute, Cambridge, MA). FeatureCounts (v. 2.0.3) was used to generate count matrices and multiQC was used to summarize alignment statistics and counts [59–61]. Genes with <10 maximum reads across all samples were omitted from further analysis. Differential gene expression analysis was performed using DESeq2 (v. 1.48.1) [62], hciR (v. 1.8), and hciRdata (v. 1.112) R packages. Differential gene expression significance cutoffs were set at |fold-change| ≥1.5 (|log_2_fold change|≥0.585) and p_adj_<0.05. Genes were grouped by significant differential expression changes in response to interacting effect due to siRNAs and cell cycle state (i.e. siRNA treatment affects genes differently in interphase vs. cytokinetic cells, Figure 4C), cell state only (i.e. significant differences in cytokinesis only or interphase only cell populations, comparing siCon cytokinesis to siCon interphase, Figure 4 Supplement 2A), or siRNA treatment only (i.e. significant differences in siCon only or siNups only cell populations, comparing siNups interphase to siCon interphase, Figure 4 Supplement 2D). Genes classified as interactors were omitted from heatmaps and downstream analysis of cell state only and siRNA treatment only gene sets. Gene clusters were determined via the complete linkage method.

Alternative splicing of four contrasts (siNups interphase vs. siCon interphase, siNups cytokinesis vs. siCon cytokinesis, siCon cytokinesis vs. siCon interphase, siNups cytokinesis vs. siNups interphase) was measured using rMATS-turbo (v. 4.1.1) [63, 64], and the resulting output was analyzed using the maser R package (v. 1.26.0) [65]. Splice variants with <5 reads on average across all samples were omitted from further analysis. Alternative splicing significance cutoffs were set at |percent spliced in (PSI)| > 0.1 and false discovery rate (FDR) <0.05. This paper does not report original code. Gene ontology analysis was performed on clusters of interest using the EnrichR web tool [66–68]. Pathway enrichment significance was set at p_adj_<0.1.

### Statistical analysis

When comparing raw fluorescence intensity values, samples were normalized to siControl interphase MIG intensities (Figure 1C/D, Figure 1 Supplement 1BC-E), untreated interphase nuclear intensities (Figure 2G), or siControl interphase nuclear intensities (Figure 1E/F, Figure 1 Supplement 1F-H, Figure 2J, Figure 2 Supplement 1H/I/L/M), or DMSO interphase nuclear intensities (Figure 2 Supplement 1E), within each individual experiment. When comparing two samples with multiple experimental replicates, a nested t-test was performed on the average value of each experimental replicate (n=3-5).

## Data availability

RNA sequencing data used in this study will be available on the NCBI Gene Expression Omnibus. Requests for imaging and quantification data reported or any further information and resources should be directed to and will be fulfilled by the lead contact, Katharine Ullman.

## Key resources table

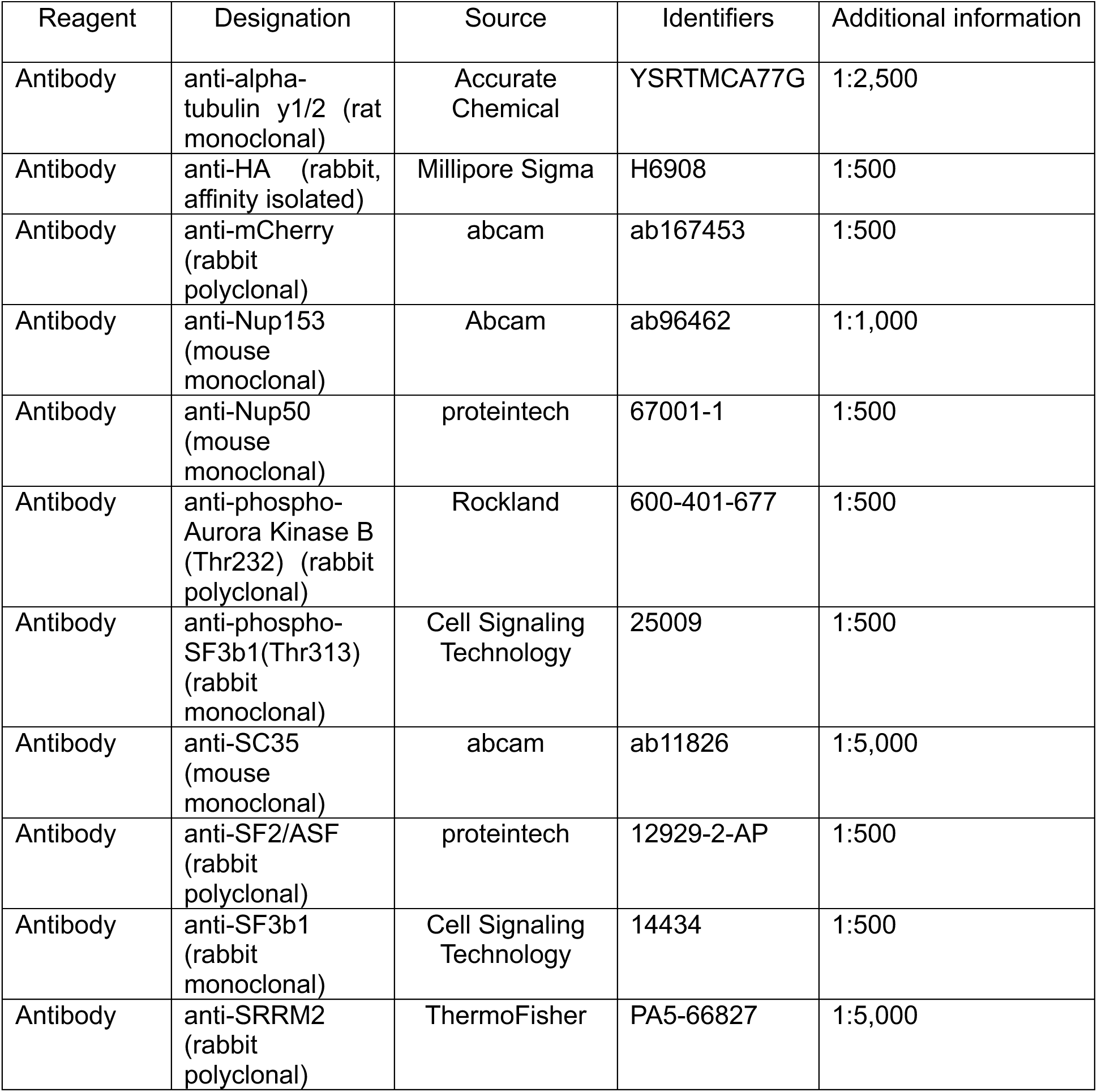
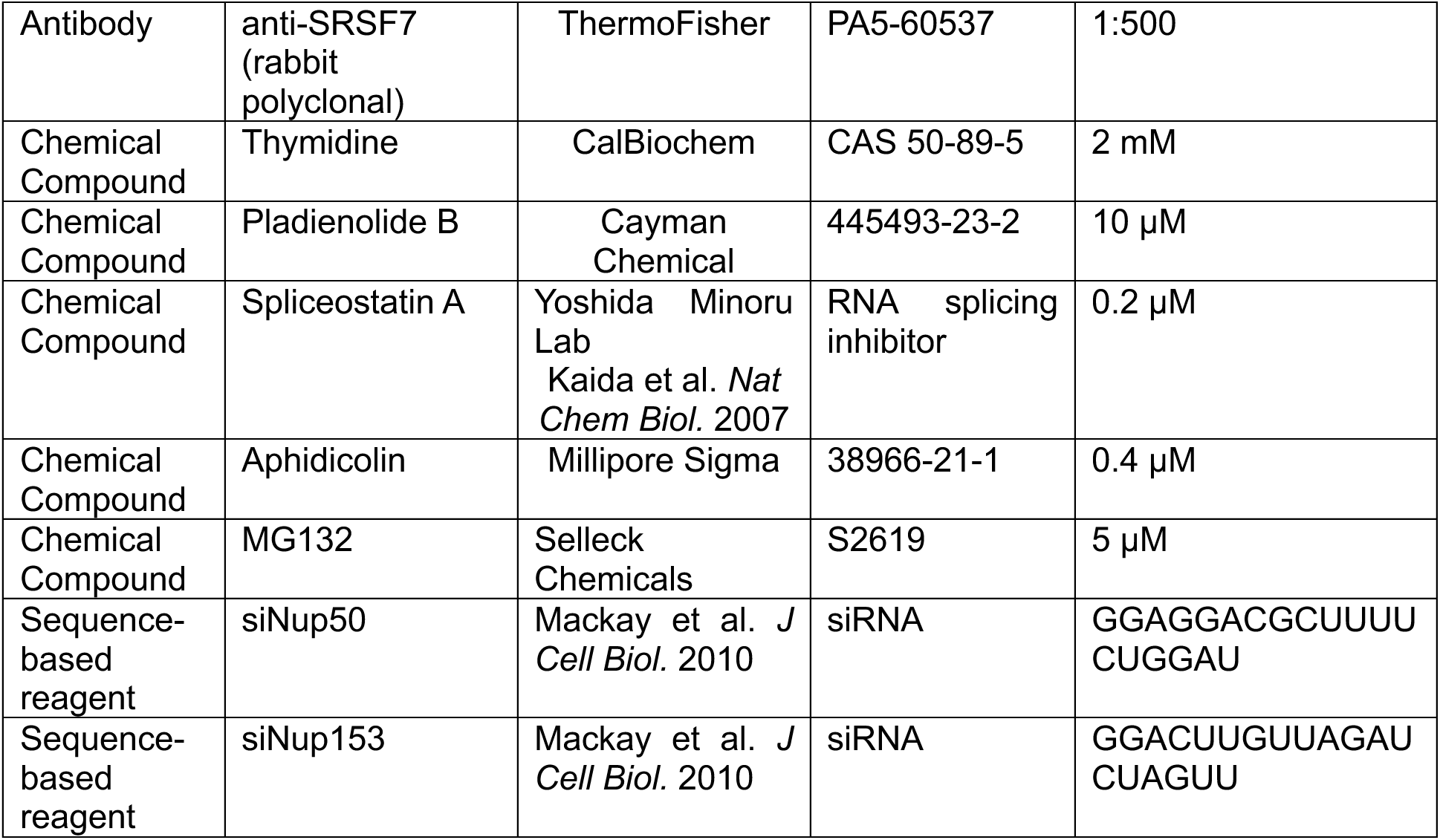

## ACKNOWLEDGEMENTS

We thank M Dasso and V Aksenova for the DLD1-SON-HA cell line. We thank M Yoshida for Spliceostatin A, L Pelkmans and A Rai for the mCherry-SRRM1 plasmid, and D Ayer for the pLVX vector. We thank L Strohacker for her guidance, expertise, and fundamental work on ACBs that this study builds upon. We thank A Turkmen for her development and optimization of the mitotic shake off procedure used to enrich cytokinetic cells. Microscopy using Leica Confocal SP8 and Nikon Automated Widefield microscopes was performed in the University of Utah Cell Imaging Core, and we thank A Classen for his imaging and microscopy expertise. Oligonucleotides were synthesized by the DNA/Peptide Facility at the University of Utah, and sequencing was performed at the DNA sequencing Core Facility at the University of Utah. We thank B Dalley, for his guidance on and knowledge of RNA sequencing techniques. Bulk RNA sequencing and analysis was performed in the High-Throughput Genomics and Cancer Bioinformatics Shared Resources at Huntsman Cancer Institute in the University of Utah, which are supported by the National Cancer Institute of the National Institutes of Health (NIH) under Award Number P30CA042014. The computational resources used were partially funded by the NIH Shared Instrumentation Grant 1S10OD021644-01A1. We also acknowledge financial support for the research reported in this publication provided by the Huntsman Cancer Foundation and the Cancer Biology and Microenvironment Program at Huntsman Cancer Institute. The content is solely the responsibility of the authors and does not necessarily represent the official views of the NIH.

## FUNDING

National Institutes of Health MPI R01 GM112080 (Wesley I. Sundquist and Katharine S. Ullman)

## AUTHOR CONTRIBUTIONS

Genevieve C, Couldwell: conceptualization, formal analysis, investigation, methodology, supervision, visualization, writing-original draft

Jessica N. Vincent-Mueller: data curation, formal analysis, investigation, methodology, software, visualization, writing – review & editing

Douglas R. Mackay: formal analysis, investigation, methodology, supervision, validation, writing - review & editing

Christopher C. Jensen: investigation, formal analysis, validation, writing -review & editing

Chris J. Stubben: data curation, formal analysis, methodology, software, validation, writing - review & editing

Li Li: data curation, software, validation, writing – review & editing

Aik Choon Tan: supervision, writing – review & editing

Wesley I. Sundquist: conceptualization, funding acquisition, methodology, project administration, supervision, writing – review & editing

Katharine S. Ullman: conceptualization, funding acquisition, methodology, project administration, supervision, writing – review & editing

## DECLARATION OF INTERESTS

The authors declare no competing interests.

---

**Figure 1 Supplement 1.**
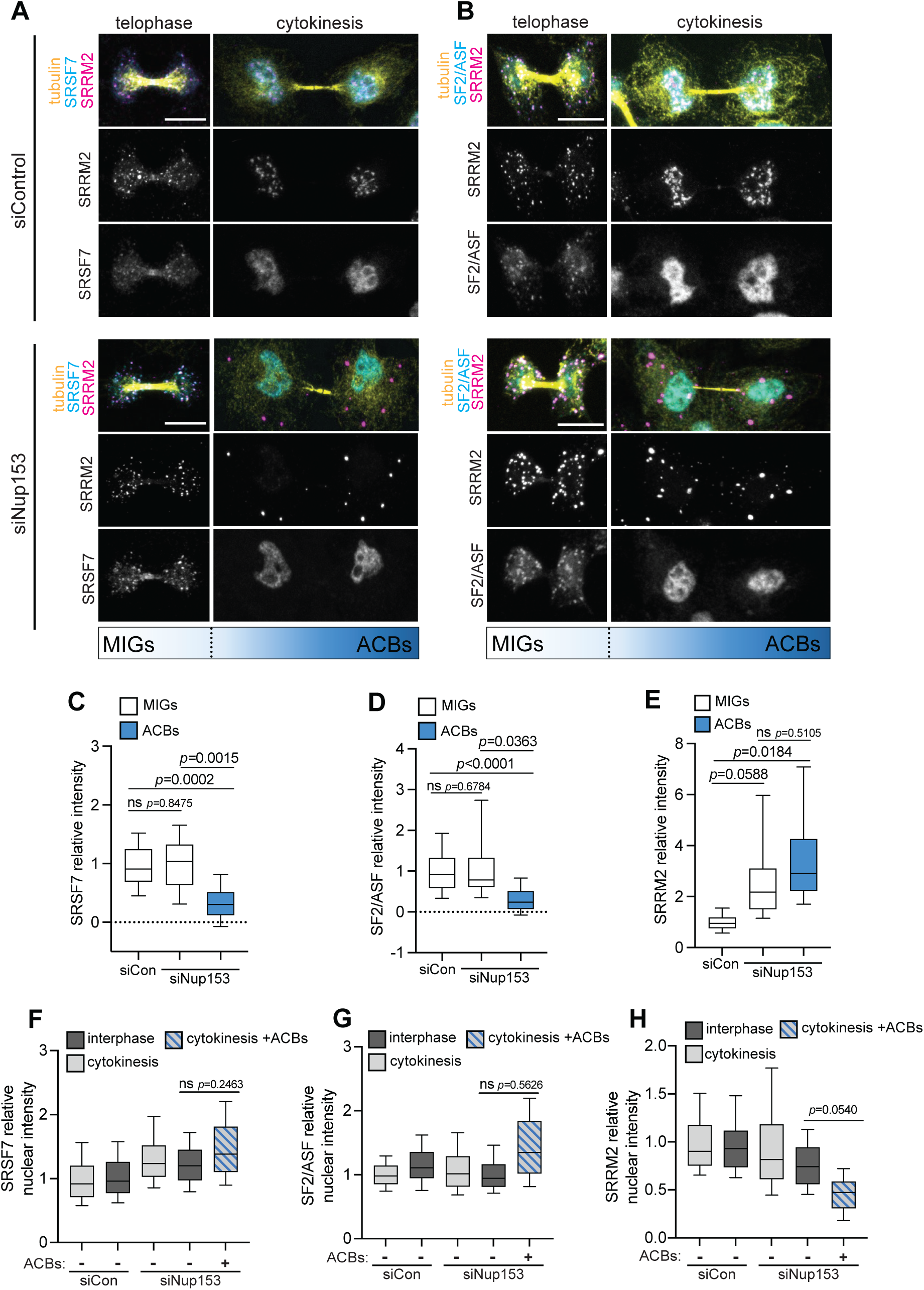
Splicing factors enrich differentially in mitotic interchromatin granules versus abscission checkpoint bodies. **A** Localization of SRSF7 and SRRM2 (mAbSC35) in confocal Z-projections of pre-permeablized HeLa cells following siCon/siNup153 treatment. **B** Localization of SF2/ASF and SRRM2 (mAbSC35) in confocal Z-projections of pre-permeablized HeLa cells following siCon/siNup153 treatment. **C, D, E** Quantification of average SRSF7, SF2/ASF and SRRM2 (mAbSC35) intensity in MIGs and ACBs, normalized to the average siCon MIG intensity. Total n≥96 cells per experimental condition collected from 4 biological replicates. **F, G, H** SRSF7, SF2/ASF and SRRM2 (mAbSC35) nuclear intensity normalized to average siCon interphase nuclear intensity. Total n≥215 cells per experimental condition collected from 4 biological replicates.

**Figure 2 Supplement 1.**
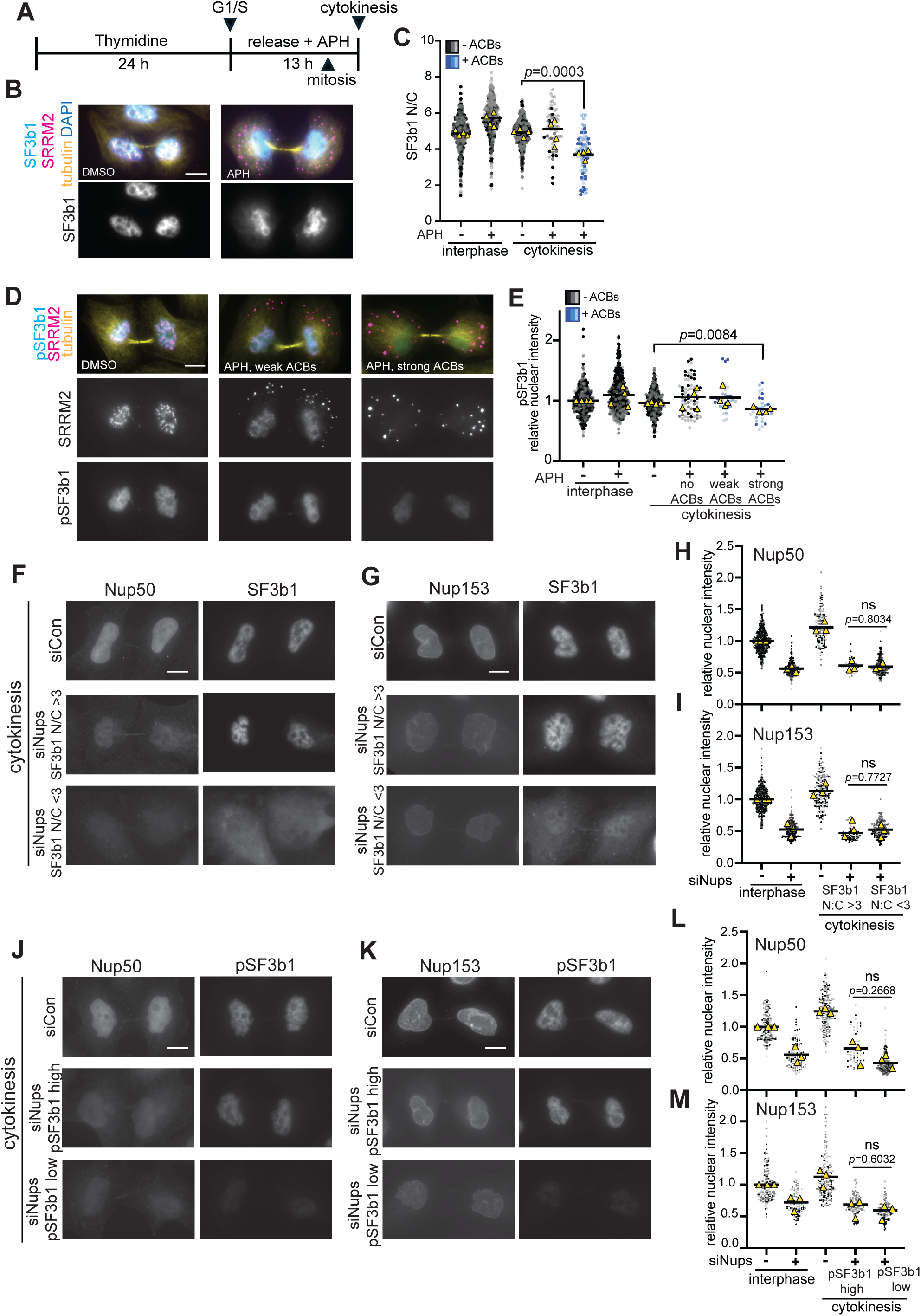
SF3b1 localization and activity are altered when ACBs are present. **A** Treatment time course of HeLa cells subjected to replication stress by treatment with 0.4 µM Aphidicolin (APH), with the peaks of cells in mitosis and cytokinesis indicated. **B** SF3b1 and SRRM2 (mAbSC35) in immunofluorescence images of cytokinetic HeLa cells treated with APH or DMSO as in (A). **C** Quantification of N/C intensity of SF3b1 in interphase and cytokinetic cells treated with APH or DMSO. Total n≥146 cells per experimental condition collected from 4 biological replicates. **D** pSF3b1 and SRRM2 (mAbSC35) in immunofluorescence images of cytokinetic HeLa cells treated with APH or DMSO as in (A). **E** Quantification of nuclear intensity of pSF3b1 in interphase and cytokinetic cells treated with APH or DMSO normalized to DMSO interphase average. ACB phenotypes were binned into “weak” and “strong” wherein a weak phenotype corresponded to SRRM2 signal in the nucleus and a strong phenotype corresponded to minimal SRRM2 in the nucleus. Total n≥124 cells per experimental condition collected from 4 biological replicates. **F, G** SF3b1, Nup50, and Nup153 in immunofluorescence images of cytokinetic HeLa cells treated with siControl or siNups. **H, I** Quantification of Nup50 (H) and Nup153 (I) nuclear intensity per cell normalized to average siCon interphase Nup50/Nup153 nuclear intensity. Total n≥154 cells per experimental condition collected from 3 biological replicates. **J, K** Immunofluorescence images of cytokinetic HeLa cells thymidine synchronized and then released for 12 h (as shown in Figure 2H) treated with siControl or siNups, with pSF3b1, Nup50, and Nup153 detected. **L, M** Quantification of Nup50 (L) and Nup153 (M) nuclear intensity per cell normalized to average siCon interphase Nup50/Nup153 nuclear intensity. Total n≥93 cells per experimental condition collected from 3 biological replicates.

**Figure 4 Supplement 1.**
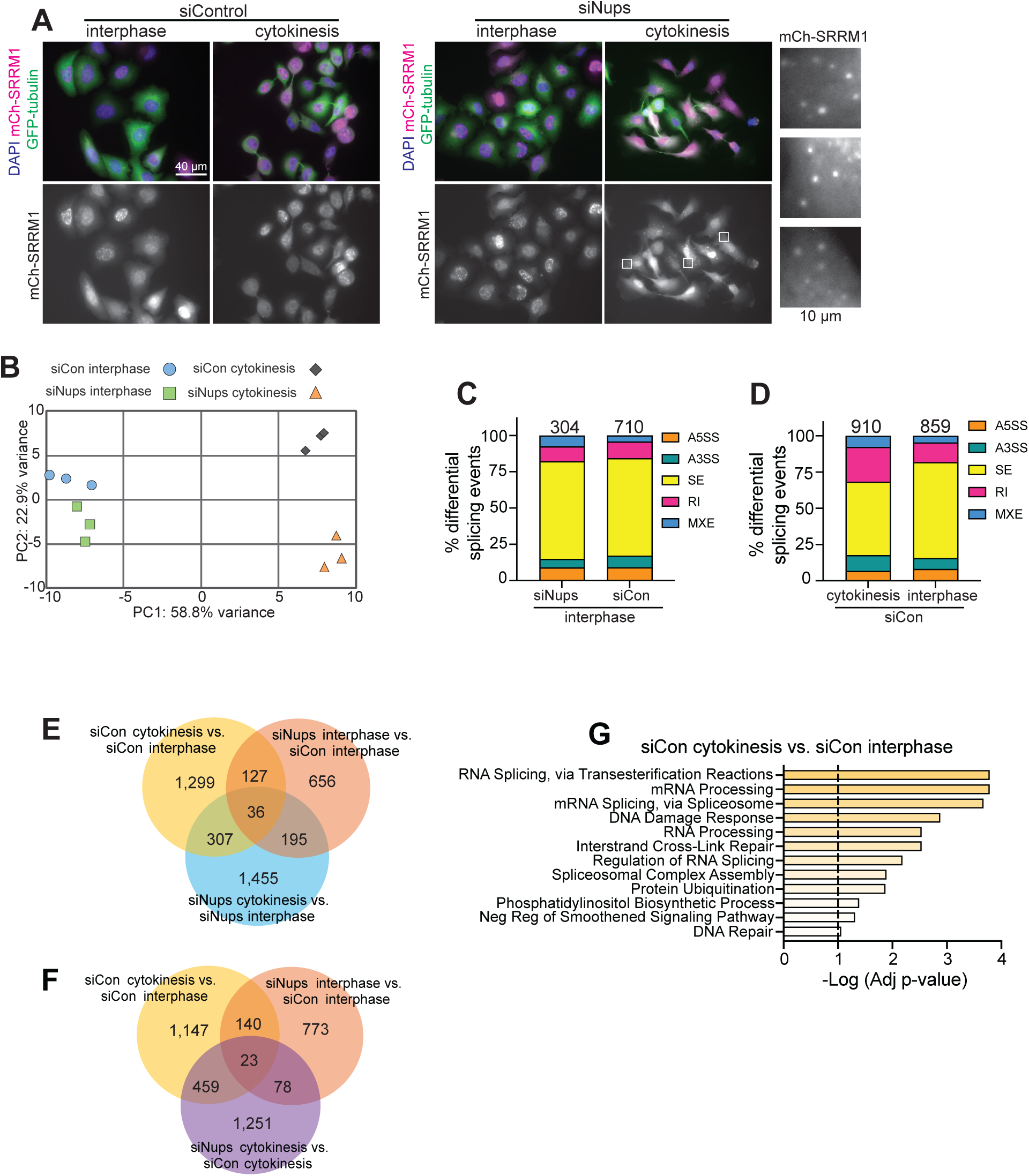
Gene expression and RNA splicing are altered when the abscission checkpoint is active. **A** Direct fluorescence detection of GFP-tubulin and mCherry-SRRM1 in interphase and cytokinetic cells, treated as in Figure 4A. Three cytoplasmic areas of siNups cytokinetic cells are enlarged at right, showing ACBs. **B** Principal component analysis of three biological replicates of each cell population sequenced. **C** Proportion of significant (FDR<0.05) alternative splicing events with |PSI|≥0.1 between siNups interphase and siCon interphase cells. Numerical values represent each differential splicing event from the comparison that occurs in either condition. **D** Proportion of significant alternative splicing events with |PSI|≥0.1 between siCon cytokinesis and siCon interphase cells. Numerical values represent each differential splicing event from the comparison that occurs in either condition. **E** Significant alternative splicing events (FDR<0.05, |PSI|≥0.1) that are identified in siCon cytokinesis vs siCon interphase (yellow), siNups interphase vs siCon interphase (orange), and siNups cytokinesis vs. siNups interphase (light blue). **F** Significant alternative splicing events (FDR<0.05, |PSI|≥0.1) that are identified in the siNups cytokinesis vs. siCon cytokinesis comparison (purple). Overlaps between this comparison and siCon cytokinesis vs siCon interphase (yellow) and siNups interphase vs siCon interphase (orange) are shown as depicted in E. **G** Biological processes gene ontologies that are significantly enriched (p_adj_<0.1) in transcripts that are alternatively spliced in the comparison between siCon cytokinesis and siCon interphase cells (1,769 transcripts, E/F, yellow circle).

**Figure 4 Supplement 2.**
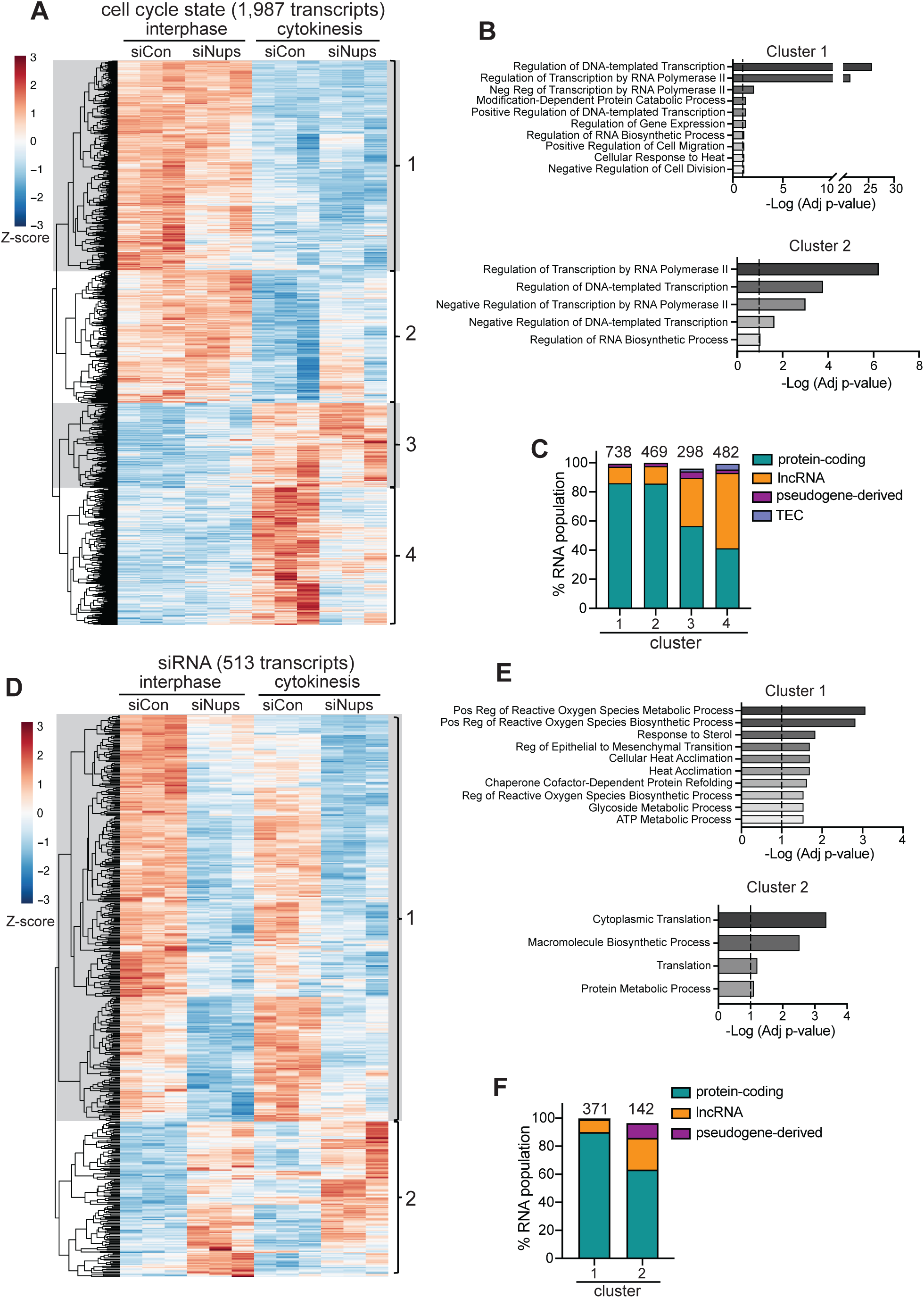
Gene expression and RNA splicing are altered under the abscission checkpoint in cell populations enriched for cytokinetic cells. **A** Heatmap and dendogram of 1,987 transcripts that had changes in expression due to cell cycle state independent of siRNA treatment (|fold change (cytokinesis/interphase)|≥1.5, p_adj_<0.05). Z-scores reflect standard deviations above (positive, red) or below (negative, blue) average regularized log-transformed read counts across samples and are scaled by row. **B** Biological process gene ontologies significantly enriched (p_adj_<0.1) in clusters 1 (top 10) and 2. Clusters 3 and 4 showed no significant enrichments. Dashed line represents p_adj_=0.1. **C** The proportion of RNA biotypes identified in (A), clusters 1-4. Biotypes with <2% representation were omitted. Numerical values above bars represent the number of transcripts in each cluster. **D** Heatmap of 513 transcripts that had changes in expression attributed to siRNA treatment only (|fold-change (siNups/siCon)|≥1.5, p_adj_=0.05). Z-scores are depicted as in A. **E** Biological processes gene ontologies significantly enriched (p_adj_<0.1) in clusters 1 (top 10) and 2. Dashed line represents p_adj_=0.1. **F** The proportion of RNA biotypes identified in D, clusters 1 and 2. Biotypes with <2% representation were omitted. Numerical values above bars represent the number of transcripts in each cluster.

## Notes

### Competing Interest Statement

The authors have declared no competing interest.

